# Identification of the toxic 6mer seed consensus in human cancer cells

**DOI:** 10.1101/2020.12.22.424040

**Authors:** Monal Patel, Elizabeth T. Bartom, Bidur Paudel, Masha Kocherginsky, Kaitlyn L. O’Shea, Andrea E. Murmann, Marcus E. Peter

**Affiliations:** Department of Medicine/Division Hematology/Oncology, Northwestern University, Chicago, IL; Department of Biochemistry and Molecular Genetics, Northwestern University, Chicago, IL; Department of Preventive Medicine/Division of Biostatistics, Feinberg School of Medicine, Northwestern University, Chicago, IL

**Author notes:** Corresponding author: Marcus Peter.

**Keywords:** DISE, cancer, 6mer seed toxicity, RNAi

## Abstract

6mer seed toxicity is a novel anti-cancer mechanism that kills cancer cells by triggering death induced by survival gene elimination (DISE). It is based on si- or shRNAs with a specific G-rich nucleotide composition in position 2-7 of their guide strand. An arrayed screen of 4096 6mer seeds on two human and two mouse cell lines identified a consensus GGGGGC as the most toxic seed. After testing two more cell lines, one human and one mouse, we found that the GGGGGC seed while also toxic to murine cells, is more toxic to human cells, suggesting that the evolution to use of Gs as part of the toxic seeds is still slowly evolving, with Gs more common in the human toxic seeds. While new RNA Seq and bioinformatics analyses suggest that the GGGGGC seed is toxic to cancer cells by targeting GCCCCC seed matches in the 3’ UTR of a set of genes critical for cell survival, we now directly confirm this by identifying a number of genes targeted by this seed. Furthermore, by using a luciferase reporter fused to the 3’ UTR of these genes we confirm direct and specific on-targeting of GCCCCC seed matches. Targeting is strongly attenuated after mutating the GCCCCC seed matches in these 3’ UTRs. Our data confirm that an siRNA containing the GGGGGC seed kills cancer cells through its miRNA like activity and points at artificial miRNAs, si- or shRNAs containing this seed as a potential new cancer therapeutics.

## Introduction

micro(mi)RNAs are small noncoding double stranded RNAs that negatively regulate gene expression at the post-transcriptional level. Activity of miRNAs is determined by the seed sequence (position 2-7/8) in the guide strand of the miRNA (1, 2). The guide strand is incorporated into the RNA induce silencing complex (RISC), where it engages the targeted mRNA by binding to seed matches located mostly in the 3’ untranslated region (UTR) that are complementary to the miRNA’s seed (3). The human genome is estimated to contain ~8300 miRNAs (4). They are predominantly regulating differentiation and development, and have been shown to be deregulated in virtually all human cancers, where they can function as tumor suppressors or oncogenes (oncomiRs) (5–7).

We previously discovered that a large number of si-and shRNAs were toxic to all cancer cells independent of hitting their intended target (8), in a process believed to be a special form of off-target effect of RNA interference (RNAi) (9). It involves simultaneous activation of multiple cell death pathways (8) and cancer cells cannot become resistant to this form of cell death. For an si- or shRNA to be toxic, a 6mer seed is sufficient (10). We demonstrated that Ago2, a critical component of the RISC, was required for cells to die through 6mer seed toxicity (6mer Seed Tox), and that the toxic si/shRNAs acted like miRNAs by targeting the 3’ UTRs of essential survival genes (10). The resulting form of cell death was therefore called DISE (for death induced by survival gene elimination). DISE was recently confirmed in the context of prostate cancer (11). An arrayed screen of all possible 4096 6mer seeds in a neutral 19mer siRNA on two human and two mouse cell lines identified the most toxic seeds and a strong conservation across tissues and species. The most toxic 6mer seeds were high in G nucleotides at the 5’ end of the seed (12). Consistent with a previous study that reported that most highly expressed genes regulating cell survival are devoid of seed matches for major miRNAs (13), we found that survival genes are significantly enriched in C-rich seed matches in their 3’ UTR (12). This suggested that while in the context of si/shRNAs that are designed to selectively silence single genes DISE could be viewed as an off-target effect, in the context of the biological function of miRNAs, this activity should be considered an on-target activity. In agreement with this interpretation, are findings that established major tumor suppressive miRNA families such as miR-34/449-5p and miR-15/16-5p kill cancer cells largely through 6mer Seed Tox by targeting hundreds of survival genes (12, 14, 15). si/shRNAs with a toxic 6mer seed can be used to treat cancer in mice. We demonstrated this in xenografted ovarian cancer without eliciting damage to normal tissues (16).

The most consistently toxic 6mer seed consensus we described was GGGGGC (12). We now demonstrate that an siRNA containing this consensus toxic seed motif (siGGGGGC) kills cancer cells indeed by targeting GCCCCC seed matches in the 3’ UTR of abundant mRNAs, many of which code for essential survival genes. Using an eCDF plot, we show a significant shift in the downregulated genes containing GCCCCC seed matches. In addition, we identify the most downregulated genes, many of which contain multiple GCCCCC seed matches in their 3’ UTR and demonstrate that silencing the ten most downregulated and highly expressed genes using an siRNA SmartPool induces cell death in transfected cells. Finally, we have mutated GCCCCC seed matches in selected targeted genes and show that they cannot be efficiently silenced anymore by siGGGGGC when their 3’ UTRs were fused to a luciferase reporter gene. These data strongly suggest that 6mer Seed Tox is an on-target mechanism regulating cell fate. When comparing the data of the siRNA screen between two human and two murine cell lines, we noticed a subtle difference in the composition of the toxic 6mer seed consensus (6merdb.org). We have now screened two more cell lines, one human and one mouse, and found that while siRNAs with a high G content are indeed toxic to both human and mouse cell lines, the three mouse cells are slightly less susceptible to high G containing siRNAs when compared to the three human cell lines. This points at a subtle evolutionary difference between human and mouse cells that should be considered when testing toxic 6mer seed containing si/shRNAs such as siGGGGGC in preclinical mouse models.

## Results and discussion

### The most toxic 6mer seeds differ slightly between human and mouse

Our recent arrayed screens of siRNAs containing all 4096 6mer seeds embedded in a neutral 19mer siRNA in two human cancer cell lines (HeyA8, ovarian and H460, lung cancer) as well as in two murine cell lines (M565, liver and 3LL, lung cancer) suggested that irrespective of tissue of origin or species, the most toxic 6mer seeds were rich in G, particularly at the 5’ end of the seed (12). Minor differences were seen between the two human and two mouse cell lines. To determine whether this holds when more cell lines representing additional tissues are tested, we screened all 4096 siRNAs on two more cell lines, both with brain as the tissue of origin: the human neuroglioma cell line H4 and the murine glioma cell line GL261 (**Fig. 1A**). When plotting the average nucleotide composition of the 6mer seed of the 100 most toxic and 100 least toxic seeds of both cell lines, the results appeared to be very similar to the ones we had obtained previously for the other four cell lines. Again, the most toxic seeds were high in Gs at the 5’ end of the seed, and the least toxic seeds were AU-rich with an A or U in the first two seed positions (**Fig. 1B**). The similarities between cell lines was also seen in a Pearson correlation analysis of the entire screen comparing each of the glioma cell line results with the average of the other two cell lines from the same species (**Fig. 1C**). However, when assessing the most toxic 6mer seeds for all 6 cell lines, subtle differences between human and mouse cells became apparent. The average seed composition of the most toxic seeds was more strongly dominated by Gs in the three human cells compared to the three mouse cell lines (**Fig. 1D, 1E**). These sequence logos (generated with Weblogo) are a graphical representation of the relative frequency of each nucleotide in each of the 6 positions of the 6mer seed. Existing sequence logo methods only permit a visual representation. We therefore developed a framework for statistical analysis of 6mer toxicity data using multinomial mixed effects regression models. This approach allows us to compare differences in sequence patterns between groups and positions, and provides both a statistical testing framework for comparing nucleotide composition between groups and positions, as well as estimates of the differences. Applying this new approach we found that there were statistically significant differences in nucleotide patterns between human and mouse composition of the top 100 most toxic seeds (p<0.0001) (**Fig. 1E**). Compared to G, nucleotides A (odds ratio OR=2.49, p<0.0001), C (OR=1.96; p<.0001) and U (OR=1.51; p=0.020) were more likely to occur in mice than in humans. The differences between groups did not vary by position (p=0.245, group × position interaction term). The difference in toxic seed composition was also visible when ranking all 4096 seeds according to the highest average toxicity in the human cells (note, the different colors between the three human and the three mouse cell lines in **Table S1**). The siRNA harboring the consensus GGGGGC was the 8th most toxic seed for human cells but only the 110th most toxic seed for the three mouse cells (marked in a red font in **Table S1**). These data have now been added to the 6merdb.org site. They suggest that while G-rich 6mer seeds are highly toxic to both human and mouse cells, the very high toxicity of G rich seeds is most prominent in human cells. In human cells the average toxic seed consensus is GGGGGC.

**Figure 1.**
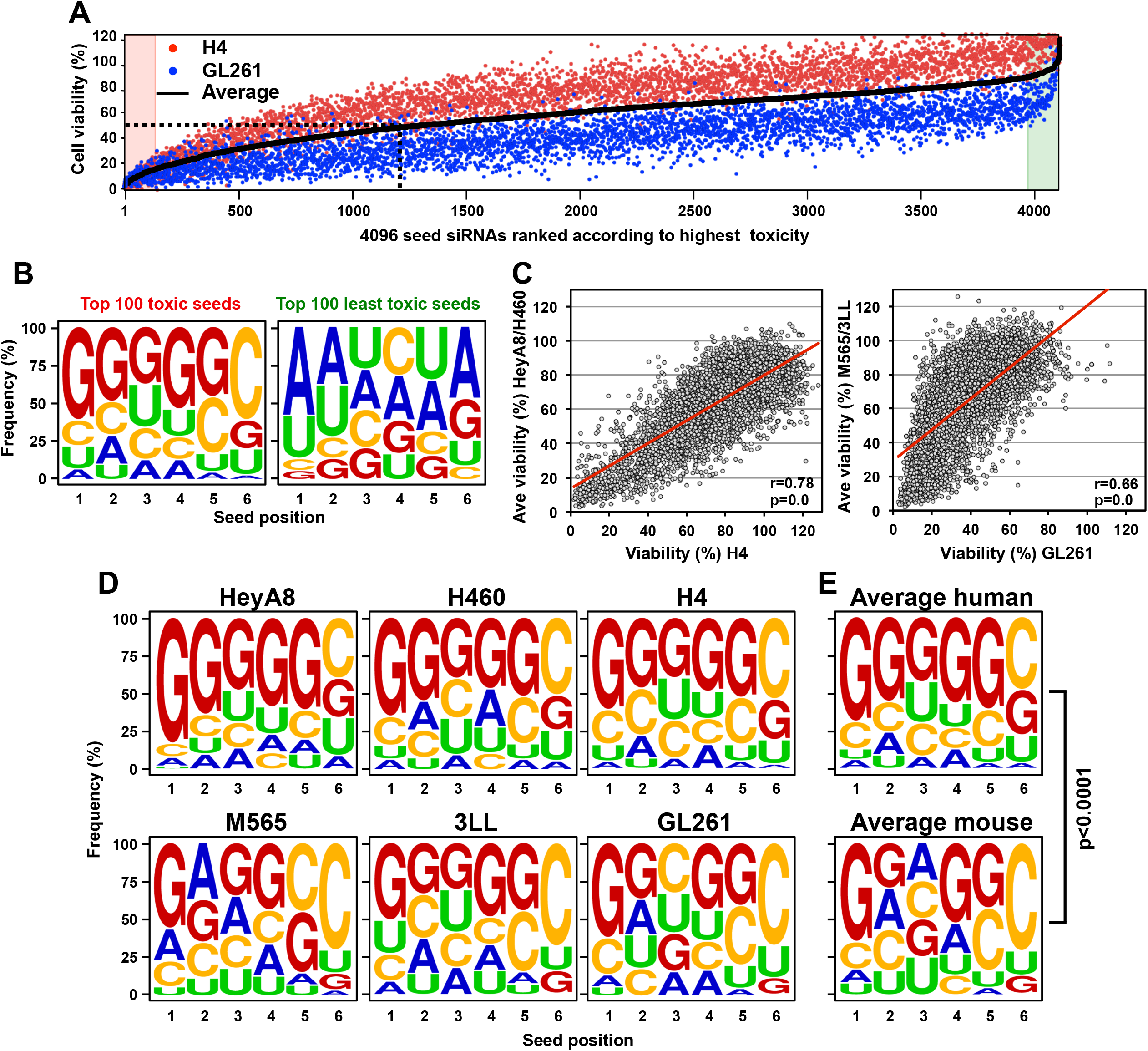
Subtle difference in the composition of the most toxic 6mer seed between human and mouse cancer cell lines. (A) *Top:* Results of the 4096 6mer seed duplex screen in a human (H4) and a mouse (GL261) glioma cell line. Cells were reverse transfected in triplicates in 384 well plates with 10 nM of individual siRNAs. The cell viability of each 6mer seed duplex was determined by quantifying cellular ATP content 96 hrs after transfection. All 4096 6mer seeds are ranked by the average effect on cell viability of the two cell lines from the most toxic (left) to the least toxic (right). We consider an siRNA highly toxic if it reduces cell viability 90% or more and moderately toxic if it reduces cell viability 50% or more (black stippled line). (B) Average seed composition of the 100 most (top) and 100 least (bottom) toxic 6mer seeds in H4 and GL261 cells screened with 4096 6mer seed containing siRNAs. (C) *Left:* Regression analysis showing correlation between the 6mer Seed Tox observed in H4 cells and the average of the two previously tested human cell lines HeyA8 and H460. *Right:* Regression analysis showing correlation between the 6mer Seed Tox observed in GL261 cells and the average of the two previously tested murine cell lines 3LL and M565. p-values were calculated using Pearson correlation analysis. (D) Seed composition of the 100 most toxic 6mer seeds in human (top) and three murine (bottom) cell lines. (E) Average most toxic seed composition of all human (top) and all murine (bottom) cell lines. P-value is derived from a multinomial mixed effects regression model.

### siGGGGGC is highly toxic to multiple cancer cells and preferentially kills cancer cells that cannot properly deal with intracellular stress

We recently reported that the prototypical tumor suppressor miRNA, miR-34a-5p, is toxic to various cancer cells mostly through 6mer Seed Tox by targeting the 3’ UTR of multiple genes that are critical for the cell survival (12). miR-34a-5p carries two Gs as the first two nucleotides in its 6mer seed GGCAGU, consistent with the observation of a high G content towards the 5’ end of the most toxic seeds. We now compare the potency of siGGCAGU to siGGGGGC, the seed of which is present among other miRNAs, in miR-1237-5p. Both siRNAs had a chemically modified passenger strand to prevent loading into the RISC (12, 17). When transfected into the OC cell line HeyA8, both siRNAs strongly slowed down cell growth (**Fig. 2A**) and reduced cell viability compared to the nontoxic siNT1, regardless of whether the siRNAs were introduced into cells by forward or reverse transfection (**Fig. 2B**). Morphologically, cell death was very similar to what we had reported for toxic 6mer seed containing CD95L derived siRNAs that induced multiple cell death pathways with a dominant necrotic component (**Fig. 2C**) (8). Cell death was not cell specific as siGGGGGC was also toxic to other human cancer cell lines (**Fig. 2D**, and (18)).

**Figure 2.**
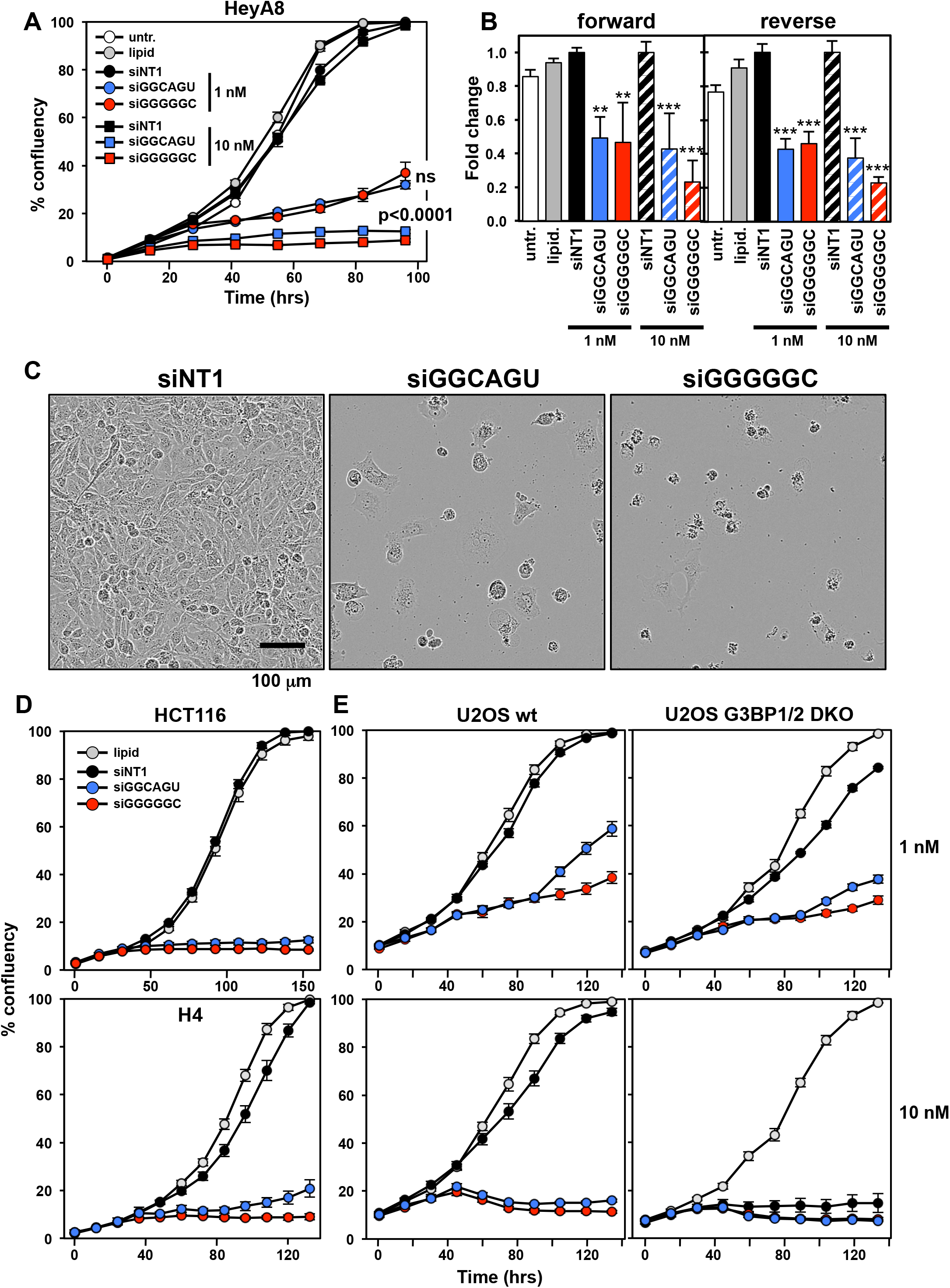
Both siGGCAGU and siGGGGGC are toxic to cancer cells. (A) Change in confluence (growth) over time of HeyA8 cells reverse transfected with either 1 or 10 nM of siNT1, siGGCAGU, or siGGGGGC. ANOVA p-values between the two toxic siRNAs are shown. ns, not significant. Each data point represents mean ± SE of six replicates. (B) Percent viability (ATP content) 96 hrs after transfecting HeyA8 cells (forward or reverse) with 1 or 10nM of the indicated siRNAs, or with lipid only or left untreated. Experiment was done in 6 replicates. *** p<0.0001, ** p<0.001. (C) Phase contrast images showing morphology of HeyA8 cells reverse transfected in A at 10 nM 4 days post transfection. (D) Change in confluence over time of HCT116 (top) and H4 (bottom) cells reverse transfected with 10 nM of siNT1, siGGCAGU, or siGGGGGC. Each data point represents mean ± SE of six or four replicates, respectively. (E) Change in confluence over time of U2OS wt and U2OS G3BP1/2 DKO cells reverse transfected with 1 nM (top) or 10 nM (bottom) of siNT1, siGGCAGU, or siGGGGGC. Each data point represents mean ± SE of three replicates.

Our previous data suggested that DISE is causing a strong stress response in cells resulting in ROS production and DNA damage (8). Cells show signs of severe stress with large intracellular vesicles forming before the onset of cell death (8) (**Movies S1-S3**). Two types of organelles in the cytosol typically form as a consequence of stress and altered RNA homeostasis (stress granules, SGs and P-bodies, PBs) (19). Recently, G3BP1 and G3BP2, two RNA binding proteins were shown to be essential for cells to form stress granules (SGs) (20–22), but not for the formation of P-bodies (20). We have now tested U2OS G3BP1/2 double knock out (DKO) cells (22) and found that the DKO cells are as sensitive to both siGGCAGU and siGGGGGC as wt cells when transfected at 1 nM (**Fig. 2E**, top panels). However, when transfected with 10 nM siRNAs, the DKO cells even died in response to the control siRNA siNT1 (**Fig. 2E**, bottom panels). These data suggest that cells that cannot properly respond to stress are hypersensitive to RNAi mediated cell death. It is known that cancer cells experience increased intracellular stress. An increased susceptibility of stressed cells to DISE could be another reason for why *in vivo* only cancer cells seem to die from DISE (16).

### siGGGGGC induced cell death is similar to the one induced by siGGCAGU and carboplatin

To test how siGGGGGC induces cell death, we performed an RNA Seq analysis of HeyA8 cells at both 24 and 48 hrs after transfection with 10 nM of siNT1 or siGGGGGC. The genes downregulated in the siGGGGGC transfected cells, were enriched in a reference set of recently defined survival genes (10) (**Fig. 3A**), similar to the activity we reported for the two tumor suppressive miRNAs miR-34a-5p and miR-15/16-3p and their matching 6mer seed containing siRNAs (12, 14). A gene ontology analysis suggested that the form of cell death induced by siGGGGGC, siGGCAGU (the miR-34a-5p 6mer seed), and even the chemotherapeutic drug carboplatin were very similar with shared GO terms we had previously reported to characterize DISE (10, 12, 23). To determine whether siGGGGGC targets mRNAs that contain GCCCCC 6mer seed matches in their 3’ UTR, we analyzed mRNAs deregulated in HeyA8 cells 24 hrs after transfection with siGGGGGC (**Fig. 3C**). Using a Sylamer analysis as a method to identify enriched sequences in a list of downregulated mRNAs (24), we found a strong enrichment of the GCCCCC sequence expected to be targeted by siGGGGGC in the 3’ UTR of downregulated mRNAs (**Fig. 3C**, right) but not in their ORF (**Fig. 3C**, left). An eCDF plot, an established form of analysis to identify target engagement by miRNAs, confirmed the targeting of multiple mRNAs by siGGGGGC (**Fig. 3D**); the result of a similar analysis in cells transfected with siGGCAGU is shown for comparison. This analysis suggested that siGGGGGC induced cell death by targeting GCCCCC seed matches present in the 3’ UTR of genes.

**Figure 3.**
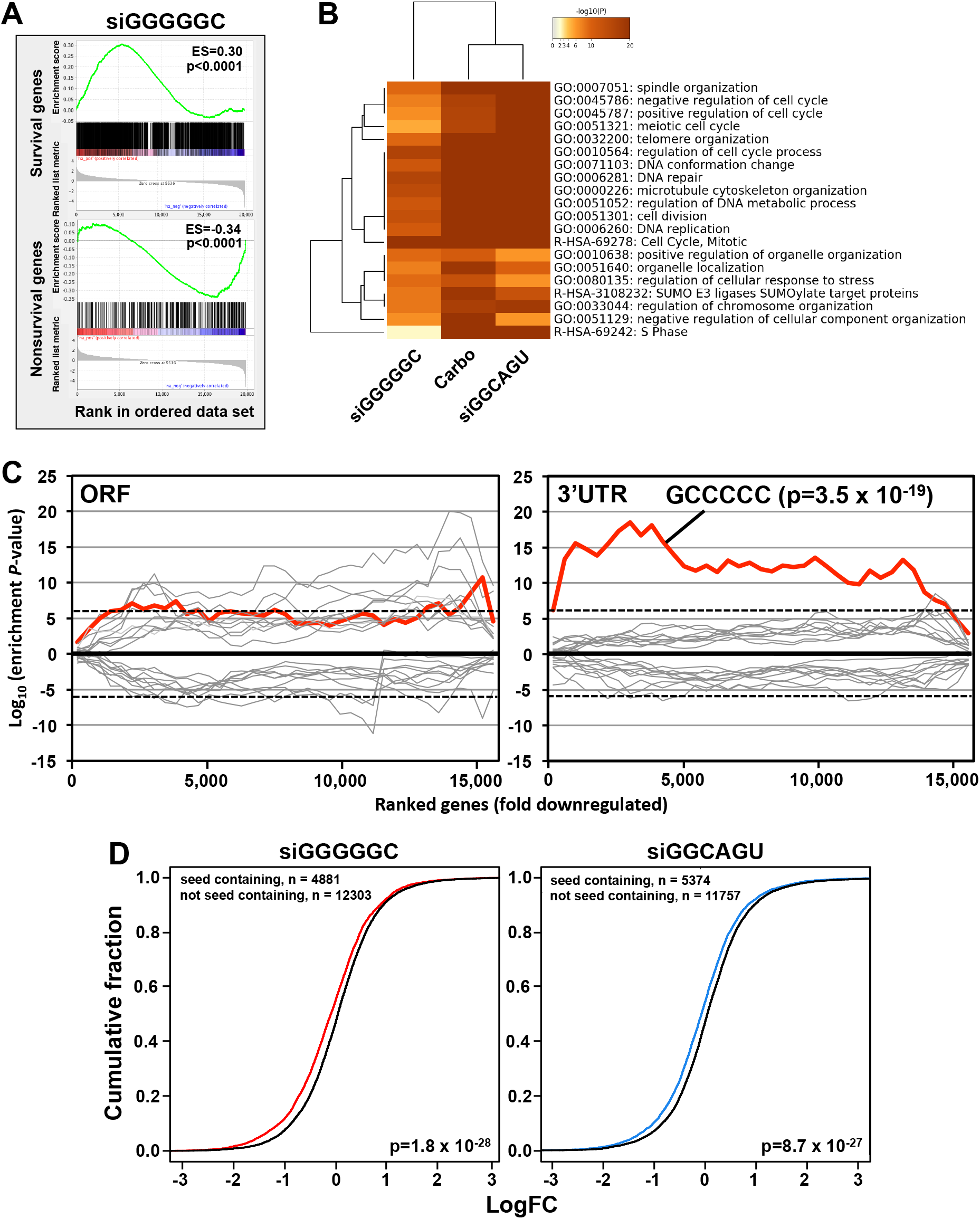
siGGGGGC is toxic to cancer cells through DISE induction. (A) Gene set enrichment analysis for a group of 1846 survival genes (top panel) and 416 non-survival genes (bottom panel) (10) 48 hrs after transfecting HeyA8 cells with siGGGGGC. siNT1 served as a control. p-values indicate significance of enrichment; the enrichment score (ES) is shown. (B) Metascape gene ontology analysis comparing genes significantly downregulated from the RNA-seq data in HeyA8 cells treated with siGGGGGC, carboplatin or siGGCAGU for 48 hrs. (C) Sylamer analysis (6mers) for the list of open reading frames (ORFs; left) and 3‘ UTRs (right) of mRNAs in cells treated with 10 nM siGGGGGC for 24 hrs sorted from down-regulated to up-regulated. Enrichment of the GCCCCC sequence is shown in red, other sequences are in grey. Bonferroni-adjusted p-value is shown. (D) eCDF plots of all mRNAs and mRNAs containing a unique 6mer seed match for either siGGCAGU (left) or siGGGGGC (right). p-values were calculated using a two-sample Kolmogorov-Smirnov (K-S) test (alternative hypothesis= “greater”).

### siGGGGGC acts like a miRNA by targeting GCCCCC seed matches enriched in the 3’ UTR of a number of survival genes

To determine whether GCCCCC seed matches were indeed being targeted by siGGGGGC in a miRNA-like fashion, we identified the 10 most significantly downregulated and highly (base mean >1000) expressed mRNAs that contain at least one putative GCCCCC sequence in their 3’ UTR (**Fig. 4A** and **Table S2**). Of these ten genes, three were part of our curated list of survival genes (labeled red in **Fig. 4A**). Most of them contained multiple GCCCCC sequences in their 3’ UTRs (**Fig. 4B, C**). In fact, these ten most downregulated genes (out of 304 genes) contained significantly more seed matches than the ten least downregulated genes (**Fig. 4D** and **Table S2**). We confirmed by qPCR that all ten genes were effectively silenced in the cells transfected with siGGGGGC (**Fig. 4E**). When these ten mRNAs were then targeted using specific SmartPools of four siRNAs each, eight of them significantly reduced cell viability, with three of the siRNA pools being most toxic (**Fig. 4F**), two of which targeted established survival genes (PES1 and POLR2E). To confirm that siGGGGGC was indeed specifically targeting GCCCCC seed matches, we cotransfected cells with siGGGGGC and a luciferase reporter construct carrying either the wt 3’ UTR or a version with mutated GCCCCC seed matches of two of the genes we found to be important for cell survival in HeyA8 cells (PES1 and POLR2K) (**Fig. 4G**). In each case, the mutant 3’ UTR construct was more resistant to the suppressive activity of siGCCCCC than the wt construct. In summary, our data indicate that siRNAs with toxic 6mer seeds act like miRNAs by targeting 6mer seed matches located in the 3’ UTR of targeted genes and that small amounts of these siRNAs are sufficient to kill cells.

**Figure 4.**
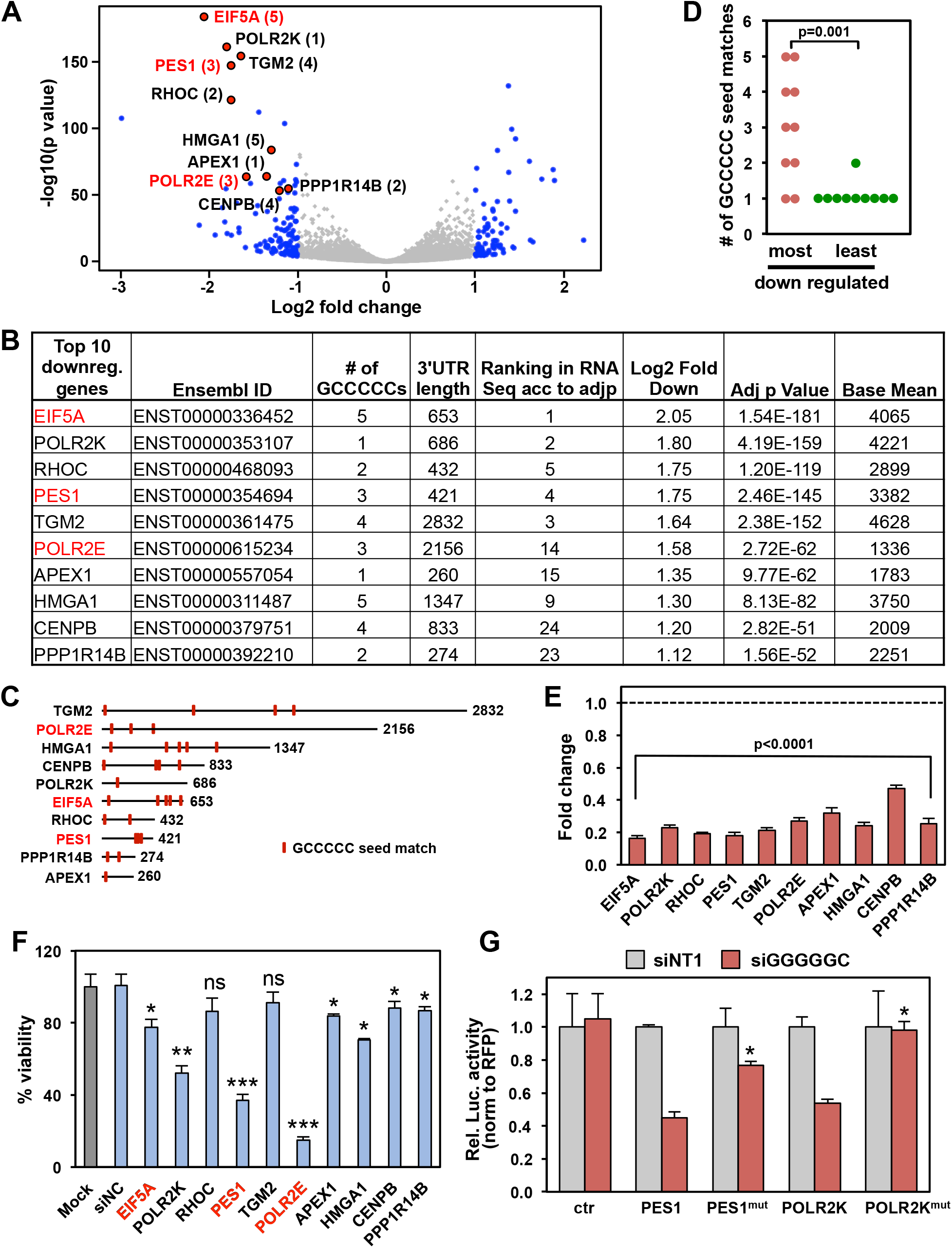
siGGGGGC kills cells by targeting GCCCCC seed matches located in the 3’ UTR of targeted mRNAs. (A) Volcano plot of deregulated genes in HeyA8 cells 24 hr after transfection with 10 nM siGGGGGC vs. siNT1. Red dots indicate genes that have one or more predicted GCCCCC seed matches in their 3’ UTR. The number of GCCCCC seed matches is given in brackets. Grey dots indicate all differentially expressed genes (with a log2 fold change <3 and >-3), blue dots indicate genes with log2 fold change>1 and FDRAdjpValue<0.05. (B) Table showing the 10 downregulated genes in A ranked (from top to bottom) according to Log2 fold change. (C) Schematic of the 3’ UTRs of the 10 genes listed in B with the location of the GCCCCC seed matches. (D) The number of predicted seed matches in the ten most and least downregulated and highly expressed genes part of the 5957 genes in the human genome that contain at least one such seed match in their 3’ UTR. p-value was calculated by Student’s T-test. (E) Real-time PCR validation of downregulation of the ten genes in A subjected to RNA seq analysis (24 hr time point). Gene expression levels in HeyA8 cells transfected with siGGGGGC were normalized to that in siNT1 transfected cells (stippled line). Each bar represents ± SD of three replicates. Students T-test was used to calculate p-values. (F) Viability of HeyA8 cells 96 hrs after transfection with 10 nM of siRNA SmartPools against the ten genes in A. Each bar represents ± SD of three replicates. *** p<0.0001, ** p<0.001, * p<0.05, ns, not significant. Genes that are part of our curated list of survival genes are labeled in red. (G) Relative luciferase activity of control plasmid (pmiR-Target), wild-type and mutant 3’ UTR constructs of PES1 and POLR2K. Data are shown as mean ± SD of three replicates. p-values (Students t-test, *p<0.05) represent comparison between change in relative luciferase of cells co-transfected with wt/mutant constructs and siGGGGGC normalized to cells transfected with siNT1.

6mer Seed Tox is an intrinsic anti-cancer mechanism possibly built into all genomes of multicellular organisms that harbor miRNAs. It is based on the understanding that highly expressed miRNAs must not be toxic to normal cells. Hence, 3’ UTRs of genes critical for cell survival are generally devoid of seed matches for abundant miRNAs (13). Consistent with this observation, we reported that older and more conserved miRNAs contain fewer toxic 6mer seeds and that many younger miRNAs including most miRtrons which have G-rich seeds are barely expressed in cells (12). Based on an analysis of young and old miRNAs, we concluded that the 6mer Seed Tox/DISE mechanism is at least 800 million years old. By testing two more cell lines, one human and one mouse, we now provide evidence that the G-rich toxic mechanism is still evolving and the extreme G-richness of the most toxic seeds is less pronounced in mouse cells.

We have now identified a number of direct targets of siGGGGGC which are strongly silenced in cells shortly after transfection. In principle, these targets could be used as biomarkers for silencing to confirm target engagement during a potential treatment of cancer with this toxic seed containing RNA oligonucleotide. However, there can be no definite list of targeted genes for all cancers or cancer cells, as the nature of the targeted genes will likely depend on the transcriptome of every individual cancer cell. *In vivo* targeting, however, can be established by performing RNA Seq of tumor tissue after treatment and subjecting the data to the bioinformatics analyses as presented here.

We have previously shown that siRNAs that are toxic through 6mer Seed Tox can be used to treat ovarian cancer in mice without any signs of toxicity to the mice (16). However, these experiments were done before we had identified the most toxic 6mer seeds. Using the 6mer seed of an established tumor suppressive miRNA such as miR-34a-5p, which has already been tested in a clinical trial for the treatment of various human cancers (25) may be a viable new treatment option for advanced cancers, particularly as we recently demonstrated that ovarian cancer cells resistant to platinum, the first line treatment for ovarian cancer, show an increased sensitivity to 6mer seed toxicity (18). However, rather than using the entire miRNA, our data suggest that it may be sufficient to use an artificial miRNA just containing a toxic 6mer seed. Our analysis identifies the 6mer seed GGGGGC as the consensus sequence for the most toxic 6mer seed for human cells. While there are subtle differences in the composition of the most toxic seeds for mouse cells, it should still be possible to test the GGGGGC seed containing si- or shRNAs in preclinical mouse tumor models as the GGGGGC seed is still highly toxic to mouse cancer cells.

## Materials and methods

### Cell lines

HeyA8 cells were grown in RPMI 1640 medium (Corning #10-040 CM) with 10% heat-inactivated fetal bovine serum (FBS) (Sigma-Aldrich #14009C), 1% L-glutamine (Mediatech Inc), and 1% penicillin/streptomycin (Mediatech Inc). U2OS cells deficient in G3BP1 and G3BP2 (kindly provided by Dr. Paul Taylor, St Jude’s Children’s Hospital) and 293T and H4 cells were all cultured in DMEM medium (Corning #10-013-CM) supplemented with 10% FBS and 1% penicillin/streptomycin.

### Reagents

Lipofectamine RNAimax (#13778150) & Lipofectamine 2000 (#11668019) were from Thermofisher Scientific. Optimem was purchased from Gibco (#31985-070). Cell-Titer Glo was from Promega (#G7570).

### Transfections and cell growth assessment

To assess cell growth over time using IncuCyte Zoom live-cell imaging system (Essen Bioscience), cells were either reverse or forward transfected in 96-well plates using Optimem containing optimized amounts of RNAimax and either 1 or 10nM of siRNA. For IncuCyte experiments, the following plating densities were used per 96-well: HeyA8 (1000 cells), HCT116 (2200 cells), U2OS (2000 cells). Growth curves were generated in Excel using percent confluence data over time obtained using IncuCyte Zoom software. Custom siRNA oligonucleotides were ordered from Integrated DNA Technologies and oligonucleotides were annealed according to the manufacturer’s instructions. The siRNAs containing the toxic seeds were designed as described previously (12). The sequences used were as follows:

siNT1 sense: mUmGrGrUrUrUrArCrArUrGrUrCrGrArCrUrArATT
siNT1 antisense: rUrUrArGrUrCrGrArCrArUrGrUrArArArCrCrAAA
siGGCAGU sense: mUmGrGrUrUrUrArCrArUrGrUrArCrUrGrCrCrATT
siGGCAGU antisense: rUrGrGrCrArGrUrArCrArUrGrUrArArArCrCrAAA
siGGGGGC sense: mUmGrGrUrUrUrArCrArUrGrUrGrCrCrCrCrCrATT
siGGGGGC antisense: rUrGrGrGrGrGrCrArCrArUrGrUrArArArCrCrAAA

For knocking down the ten downregulated genes in **Fig. 4F**, HeyA8 cells were reverse transfected using Optimem and 0.1 μl RNAimax per 96-well in triplicates with 10 nM of either of the following ON-TARGETplus human siRNA SmartPools purchased from Dharmacon: EIF5A (#L-015739-00-0005), PES1 (#L-009542-00-0005), POLR2E (L-004739-01-0005), HMGA1 (#L-004597-00-0005), TGM2 (#L-004971-00-0005), POLR2K (#L-011979-01-0005), RHOC (#L-008555-00-0005), APEX1 (#L-010237-00-0005), CENPB (#L-003250-00-0005), PPP1R14B (#L-026574-00-0005). The Non-targeting control pool (#D-001810-10-05) was used as a negative control.

### siRNA screens and cell viability assay

The 4096 siRNAs were described and used in an arrayed screen as described previously (11). In brief, transfection efficiency was optimized for two more cell lines (H4 and GL261) individually. RNA duplexes were first diluted with Opti-MEM to make 20 μl solution of 10 nM (for H4 cells) and 25 nM (for GL261 cells) as final concentration in a 384-well plate by Multidrop Combi. Lipofectamine RNAiMAX (Invitrogen) was diluted in Opti-MEM (5.8 μl lipid + 994.2 μl of Opti-MEM for H4, 52.6 μl lipid + 947.4 μl of Opti-MEM for GL261). After incubating at room temperature for 5–10 min, 20 μl of the diluted lipid was dispensed into each well of the plate that contains RNA duplexes. The mixture was pipetted up and down three times by PerkinElmer EP3, incubated at room temperature for at least 20 min, and then, the mixture was mixed again by PerkinElmer EP3. 10 μl of the mixture was then transferred into wells of three new plates (triplicates) using the PerkinElmer EP3. 40 μl cell suspension containing 550 H4 or 550 GL261 cells was then added to each well containing the duplex and lipid mix, which results in a final volume of 50 μl. Plates were left at room temperature for 30 min and then moved to a 37°C incubator. 96 hrs post transfection, cell viability was assayed using CellTiter-Glo quantifying cellular ATP content. 25 μl medium was removed from each well, and 20 μl CellTiter-Glo cell viability reagent was added. The plates were shaken for 5 min and incubated at room temperature for 15 min. Luminescence was then read on the BioTek Synergy Neo2. The 4096 6mer seed containing duplexes were screened in three sets for each cell line. Each set was comprised of five 384 well plates. A number of control siRNAs of known toxicity (including siNT1 and siL3, (10)) was added to each plate to compare reproducibility. All samples were set up in triplicate (on three different plates = 15 plates/set). Screen results were normalized to the average viability of the cells to siNT1 correcting the variability between sets. For the viability assays in **Fig. 2B** and **4F**, 96 hrs post transfection the medium in 96-wells was replaced with 70 μl of fresh media and 70 μl of CellTiter-Glo reagent was added on top and the luminescence was then measured as above using BioTek Cytation 5.

### RNA-seq analysis

For RNA-seq experiment 100,000 HeyA8 cells were reverse transfected with siGGGGGC, siGGCAGU or non-targeting siNT1 (siUAGUCG) duplex in multiple 6-well plates in duplicates, each well with 500 μl Optimem mix containing 1 μl RNAimax per well and siRNA at 10 nM plus 2 ml media per well. Cells were lysed the next day using Qiazol for the 24 hr time point, and to the remaining wells media was changed at 24 hr and then cells were lysed with Qiazol at 48 hr post transfection. A DNAse digestion step was included using the RNAse-free DNAse set (Qiagen #79254). Total RNA was then isolated using the miRNeasy Mini Kit (Qiagen # 74004). The quality of RNA was determined using an Agilent Bioanalyzer. The RNA library preparation and subsequent sequencing on Illumina HiSEQ4000 were done by Nu-Seq core at Northwestern University (Chicago). Paired end RNA-seq libraries were prepared using Illumina Tru-Seq Total RNA Library Prep Kit and included a RiboZero rRNA depletion Reads were trimmed with Trimmomatic v0.33 (TRAILING:30 MINLEN:20) (26) and then aligned to the hg38 human genome assembly with Tophat v2.1 (27). Exonic reads were assigned to genes using the Ensembl 78 version of the hg38 transcriptome and htseq v0.6.1 (28). Differential expression analysis was carried out using the edgeR package (29) to fit a negative binomial generalized log-linear model to the read counts for each gene.

### Data analyses

GSEA was performed using the GSEA v4.0.3 software from the Broad Institute (https://software.broadinstitute.org/gsea/). From the RNA-seq data, a ranked list was generated by sorting genes according to the Log10(fold downregulation) and this list was then used to perform GSEA using the Pre-ranked function & 1000 permutations were used. The survival and non-survival gene sets previously described (10) were used as custom gene sets. The GO enrichment analysis was performed with all genes that after alignment and normalization were at least 2 fold downregulated with an adjusted p-value of <0.05 using the software available on www.Metascape.org; default settings were used. The data set on carboplatin treated HeyA8 cells was previously published (12), accession number: GSE111379.

Sylamer analysis was performed as recently described (12). 3’UTRs or ORFs were used from Ensembl, version 76.

Volcano plot was generated using RStudio. Using the list of genes differentially expressed in HeyA8 cells transfected with siGGGGGC vs. siNT1 from the RNA-seq data, the R-script plotted all genes with a log2 fold change of <3 and >-3 on x-axis and –log10pValue on y-axis.

To identify the most significantly downregulated genes in cells transfected with siGGGGGC that carry at least one putative GCCCCC seed match in their 3’UTR, we used a data set previously described with the number and location of all 6mer motifs in the 3’ UTR of all human genes (17) (sequences extracted from ensembl.org). All genes differentially expressed (adjp<0.05, >1 Log2 fold deregulated) between cells transfected with siGGGGGC versus siNT1 with a base mean of 1000 or higher were compared to the list of genes with the GCCCCC seed matches. Transcripts ranked according to higher fold downregulation in the siGGGGGC transfected cells are shown in **Table S2**. The top most highly downregulated and bottom least downregulated transcripts were used to generate **Fig. 4D**.

### Real-time PCR

Real-time PCR was performed for the top 10 most downregulated and highly expressed genes in **Fig. 4B**. Briefly, 200 ng total RNA was used to make cDNA using the High-Capacity cDNA Reverse Transcription Kit (Applied Biosystems #4368814). The qPCR was then done using the Taqman Gene Expression Master Mix (ThermoFisher Scientific #4369016) and the following primers (human) from ThermoFisher Scientific: APEX1 (Hs00172396_m1), CENPB (Hs00374196_s1), EIF5A (Hs00744729_s1), HMGA1 (Hs00852949_g1), PES1 (Hs00897727_g1), POLR2E (Hs00267554_m1), POLR2K (Hs01562397_m1), PPP1R14B (Hs02598738_g1),RHOC (Hs00237129_m1), TGM2 (Hs01096681_m1), and GAPDH control (Hs00266705_g1). The relative expression of each target gene was normalized to the level of GAPDH. Statistical analysis was performed using Student’s t-test.

### Luciferase reporter assay

293T cells were co-transfected in triplicates in 96 well plates, 24 hr post plating, with 50 ng of each luciferase reporter clones and 20 nM of either siNT1 or siGGGGGC using Lipofectamine 2000 (Thermofisher). The control plasmid, pMirTarget (#PS100062); PES1 Human 3’UTR Clone (#SC205900, NM_014303); POLR2K Human 3’UTR Clone (#SC208896; NM_005034); Mutant PES1 (#CW306514) and POLR2K (#CW306515) 3’UTR clones were all purchased from Origene. Mutant 3’UTR plasmids were generated by synthesizing 3’ UTR sequences in which the three GCCCCC seed matches in the 3’ UTR of PES1 and one in 3’ UTR of POLR2K was each changed to TGCAAA and this altered sequence was each inserted into the pmiRTarget construct by Origene. After 48 hrs, transfection efficiency was determined by measuring Red Fluorescence Protein (RFP) expression of the luciferase reporter plasmid followed by measurement of luciferase activity after lysing cells with Bright-Glo Luciferase Assay System (Promega #E2610), both using Biotek Cytation 5 plate reader. Data are shown as relative luciferase activity normalized to RFP expression.

### Empirical cumulative distribution function (eCDF) plot

From the RNA-seq data in HeyA8 cells transfected with siGGGGGC or siGGCAGU, in order to determine if the mRNAs are regulated based on presence of GCCCCC (seed match for siGGGGGC) or ACTGCC (seed match for siGGCAGU) seeds in their 3′UTR, a custom R script was used. The R script takes as input a list of genes containing the seed and a list of genes not-containing the seed in their 3’ UTR, in each case, as well as a table of logFC expression for those genes upon siGGGGGC or siGGCAGU over-expression. This script then generates the eCDF plot for the logFC expression data for each gene set. The custom scripts and the input data files for the eCDF plots are available in Code Ocean at https://codeocean.com/capsule/9825010/tree for siGGGGGC and at https://codeocean.com/capsule/8338685/tree for siGGCAGU. Kolmogorov–Smirnov (K-S) two-sample test (alternative hypothesis = “greater”) was used to compare different probability distributions shown in the eCDF plots.

### Statistical analyses

IncuCyte experiments were performed in triplicates and the data were expressed as mean ±standard error, continuous data were compared using Student’s T-tests for two independent groups. Two-way ANOVA was performed using the Stata1C software for evaluating continuous outcomes over time, choosing one component for the treatment condition as primary interest and a second component for time as a categorical variable.

#### Multinomial mixed effects regression model to analyze 6mer seed compositions

A novel approach for the analysis of sequence logo data using a generalized linear mixed model for multinomially distributed was developed. Sequence logos (generated with Weblogo (http://weblogo.berkeley.edu/logo.cgi) are a graphical representation of the nucleotides in each of the 6 position of the 6mer seed, but they only permit a visual representation. Multinomial mixed models provide estimates of the relative differences in nucleotide composition between groups and positions as well as a statistical test of such differences. In this framework, we represent each 6mer seed as 6 “repeated measures” and assume that nucleotides follow a multinomial distribution at each position. A multinomial mixed effects model is then fitted with the nucleotide (A, C, G, U) as the outcome; group, position and their interaction as the fixed effects; and 6mer id as the random effect. Correlation between the 6 positions is accounted for using “unstructured” covariance structure. Because the toxic seeds are dominated by Gs we compare the odds of either A, U, or C to that of G in each seed position, and obtained odds ratio (OR) estimates comparing groups and positions. An interaction term allows us to determine whether the nucleotide composition differences between groups vary by position. The multinomial mixed models were fitted using PROC GLIMMIX in SAS v. 9.4.

## Supporting information

Video S1 siNT1

Video S1 siGGCAGU

Video S3 siGGGGGC

## Data availability

RNA Seq data were deposited at GEO under accession number GSExxxx.

## Acknowledgement

We would like to thank Dr. Paul Taylor for providing the G3BP1/2 DKO cell line and Dr. Siquan Chen for performing the arrayed siRNA screens. This work was funded by grant R35CA197450 to M.E.P. and P30CA060553.

## Author contributions

M.P., B.P., A.E.M performed experiments; E.T.B. performed bioinformatics analyses; M.K., K.L.O. provided biostatistics support, M.P. wrote, and M.E.P conceived and wrote the manuscript.

---

**Table S1: Results of the screen of 4096 seed containing siRNAs on three human and three murine cells**.

Cells are color-coded: Most toxic seeds are in red and least toxic ones in green.

**Table S2: Fold change of all genes substantially expressed in HeyA8 cells transfected with siGGGGGC compared to siNT1.**

Analysis of a data set from an RNA Seq analysis of HeyA8 cells transfected with 10 nM of either siNT1 or siGGGGGC. The top ten most downregulated transcripts are in green the bottom ones in yellow. Data are ranked according to highest Log2 fold downregulation.

## Notes

### Competing Interest Statement

The authors have declared no competing interest.

### Summary of Updates

Added three suppl. videos

